# Re-engineering HIV envelope glycoproteins by circular permutation

**DOI:** 10.64898/2026.11.24.768802

**Authors:** Yesenia M. Acevedo-Rivera, Pearl Akamine, Lester J. Rosario-Rodríguez, Valerie Wojna, Gloria Cuevas-Rodriguez, Jose A. Lasalde-Dominicci, Abel Baerga-Ortiz

## Abstract

The HIV envelope glycoprotein complex (Env) is the primary target for vaccine development because it elicits neutralizing antibodies. Here, we present a family of HIV Env trimers engineered through strategic circular permutation (CP) of the BG505 and C06980v0c22 strains. We evaluated their quality attributes, including expression yields, oligomeric state, glycosylation profile, antigenicity against broadly neutralizing antibodies (bNAbs), and seroreactivity against plasma from people living with HIV (PLWH). Our results demonstrate that circularly permuted Env constructs maintained native-like trimeric assembly and recognition by benchmark bNAbs targeting major sites of vulnerability. MALDI-TOF MS analysis revealed glycosylation patterns that resemble those of linear Env native trimers, with a shift toward complex-type glycans associated with differential glycan processing. Crucially, the CP constructs exhibited reduced seroreactivity relative to the linear variant (*p* < 0.05), suggesting possible masking of immunodistracting non-neutralizing epitopes. Together, these results support circular permutation as a viable strategy to refine Env immunogen designs for targeted vaccine development.

## Introduction

The HIV Envelope glycoprotein (HIV Env or Env) is the only viral protein exposed on the surface of HIV virions, where it serves as an entry fusion machine through binding to the CD4 receptor and coreceptors of target immune cells [1,2]. In nature, Env is synthesized as the precursor protein gp160 and becomes proteolytically cleaved by host furin or furin-like proteases into monomeric gp120 and gp41 subunits that form the glycoprotein trimer [3–5]. Due to its importance in viral entry, Env has emerged as the prime candidate for the development of prophylactic vaccines against HIV.

Numerous HIV vaccine trials have explored a range of immunogens, immunization regimens, and modes of administration for different versions of the HIV Env protein, from single subunits to modified soluble trimers [6–17]. The first indication that monomeric gp120 could be a suitable vaccine candidate came from observations that it could protect chimpanzees from HIV-1 infection [18]. These early results led to the RV144 clinical trial in Thailand, which was the first to demonstrate partial protection (∼31%) in human subjects using a canarypox vector encoding *env* and *gag* prime with a gp120 boost [19–21]. Efforts to increase vaccine efficacy resulted in longer trimeric Envs with additional critical epitopes for the elicitation of neutralizing antibodies (**Fig. 1**). This strategy is exemplified by the development of soluble trimer BG505 SOSIP.664 gp140, which has been the benchmark template for multiple subsequent designs (Fig. 1d; Sanders et al., 2013). Phase 1 clinical trials of BG505 SOSIPs demonstrate that the immunogens are safe and can elicit autologous tier-2 neutralizing antibodies, but lack heterologous breadth [23,24].

**Fig. 1.**
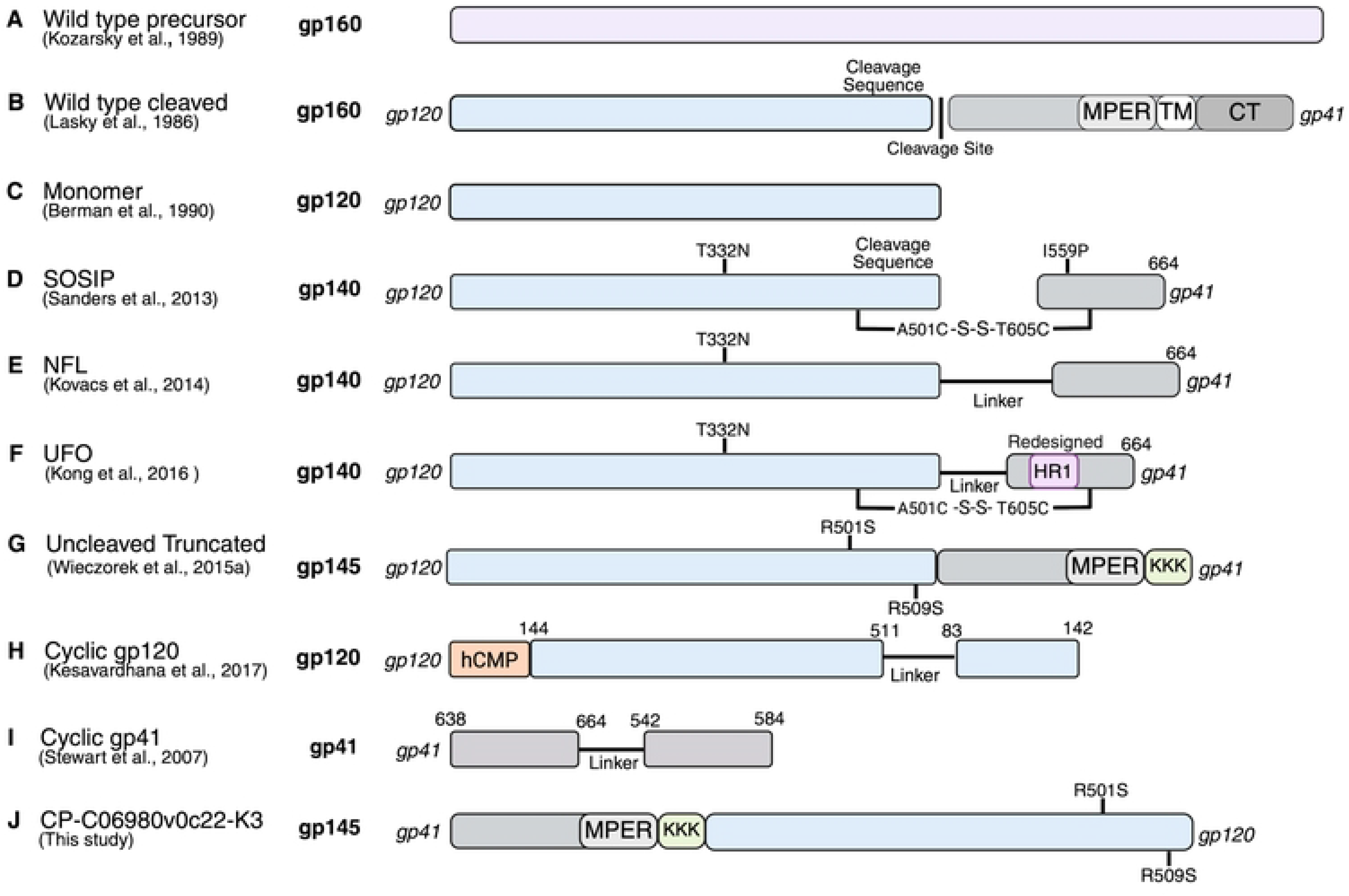
Protein engineering strategies for recombinant HIV-1 Envelope (Env) constructs. Schematic representation of the major milestones in HIV-1 Env immunogen design, tracing the progression from early monomeric subunits (gp120) to stabilized pre-fusion trimer mimics (SOSIP.664), cleavage-independent variants (NFL, UFO, Uncleaved Truncated), and cyclic monomers (cyclic gp120, cyclic gp41). Visualized engineering features include cleavage status, key stabilizing mutations (T332N, A501C, T605C, I559P, R501S, R509S), flexible linkers, C-terminal KKK motifs, coiled-coil trimerization domains, and C-terminal truncations. Building upon these approaches, this study presents CP-C06980v0c22-K3, engineered through circular permutation to introduce a new structural rearrangement. Abbreviations: CT, cytoplasmic domain of gp41; hCMP, human cartilage matrix protein; MPER, membrane-proximal external region; Redesigned HR1, redesigned heptad repeat 1; TM, transmembrane domain of gp41; NFL, native flexible linkers; and UFO, uncleaved prefusion-optimized.

Other strategies have involved re-engineered Envs containing flexible linkers that fuse gp120 and gp41 with or without SOSIP mutations [25–28]. Alternative designs produced cleavage-independent trimers that bypass the need for flexible linkers altogether by mutating the cleavage motif and truncating the membrane-proximal external region (MPER) (**Fig. 1g**). These uncleaved platforms successfully maintain quaternary integrity while reducing structural strain and show favorable biochemical profiles in early evaluations [29,30]. Significantly, this approach produced the CO6980v0c22 gp145 candidate, which successfully bridged the preclinical gap and completed Phase I clinical evaluation [31]. Yet, despite promising preclinical efficacy and safety profiles, these constructs failed to elicit favorable immunological responses in clinical trials [32–34].

Circular permutation (CP) is a protein design strategy that rearranges the order of amino acids in the primary sequence and creates new N- and C-termini [35]. CP has been used to stabilize isolated gp120 constructs into homogeneous native-like trimers that suppress immunodominant non-neutralizing decoys and selectively expose conserved sites of vulnerability (Fig. 1h; Kesavardhana et al., 2017; Saha et al., 2012). CP was also used to stabilize gp41, resulting in a protein with improved attributes for *in vitro* biophysical studies and drug discovery (**Fig. 1i**; Stewart et al., 2007). However, because these successes were confined to standalone components, extending circular permutation to the full-length Env sequence offers a mechanistically sound approach to stabilizing native trimers.

In this work, we report circularly permuted HIV-1 envelope glycoprotein gp145 (CP-Env), a reengineered uncleaved construct generated by repositioning gp41 to the N-terminus and gp120 to the C-terminus of the Env protein sequence, combined with MPER truncation and flexible linkers (**Fig. 1j**). This construct preserves the native-like trimeric architecture, maintains wild-type glycosylation patterns, and remains reactive with both broadly neutralizing and patient-derived antibodies. Overall, our results support circular permutation as a viable strategy for designing Env-derived HIV vaccine candidates.

## Materials and Methods

### Design of circularly permuted HIV envelope proteins

The genes encoding circularly permuted HIV envelope proteins (CP-Envs) were synthesized by GeneArt (Invitrogen by Thermo Fisher Scientific, Waltham, MA). All constructs were designed and cloned into separate pcDNA3.4-TOPO vectors that included a tissue plasminogen activator (tPA) signal sequence to ensure secretion into the culture medium. The constructs included the gp41 subunit with the MPER region, followed by a linker sequence of varying amino acid length connecting to the gp120 subunit. The linker sequences were **L6:** GSAGSA, **L20:** SAGSAGSSRSAGSAGSAGS, and **K3:** the MPER-stabilizing sequence KKK. Constructs CP-BG505-L6 and CP-BG505-L20 were derived from the HIV-1 BG505 Env sequence with GenBank accession number ABA61515.1 [39,40] and engineered with the T332N mutation (HXB2 numbering) to restore the potential N-linked glycosylation site (PNGS) at position 332. Constructs CP-C06980v0c22-K3 and CP-C06980v0c22 were derived from the codon-optimized C06980v0c22 *env* gene [41], which is described in detail in US Patent 10,081,658 B2 (SEQ ID NO: 20) [42]. CP-C06980v0c22-K3 includes the MPER stabilizing sequence, while CP-C06980v0c22 does not.

### Production of circularly permuted HIV envelope proteins

Each CP-Env was first expressed via transient transfection of FreeStyle™ CHO-S® cells (Invitrogen, Carlsbad, CA) using the FreeStyle™ MAX CHO Expression System (Life Technologies, Carlsbad, CA) in accordance with the manufacturer’s protocol. Briefly, cells were seeded at 1.0 x 10^^6^ cells/mL (98% cell viability) in 125 mL Erlenmeyer baffled cell culture flasks (Corning, Corning, NY) with 30 mL of serum-free FreeStyle™ CHO Expression Medium (Invitrogen, Carlsbad, CA) supplemented with 8 mM Glutamax (Fisher Scientific, Waltham, MA). DNA–lipid complexes composed of plasmid DNA and FreeStyle™ MAX Reagent (Invitrogen, Carlsbad, CA) diluted in OptiPRO™ SFM (Invitrogen, Carlsbad, CA) were prepared, incubated for 10 min at room temperature, and added to the cells. Cultures were incubated under standard conditions (37°C, 8% CO_2_, 135 rpm). Stable cell line transfection was then achieved using 400 μg/mL G418 geneticin (Gibco, Grand Island, NY) for plasmid selection and 200 μg/mL G418 geneticin for maintenance. To increase protein production, cultures were gradually scaled up to 300 mL of media in 1 L Erlenmeyer baffled cell culture flasks (Corning, Corning, NY) at 80 rpm. Samples were collected to determine cell viability with trypan blue (Bio-Rad, Hercules, CA), western blotting, and production titers. The cell suspensions were harvested after seven to nine days in culture. Cells were removed by three rounds of centrifugation at 500 x g for 10 minutes and filtered through 0.2 μm sterile PES filters (Nalgene, Thermo Fisher Scientific, Rochester, NY). The resulting supernatant containing the secreted CP-Env protein was collected and stored at −80 °C in polycarbonate storage bottles (Corning, Corning, NY) until further processing.

### Production titers

We quantified titers from culture supernatants by a reversed-phase HPLC assay previously described by our group [43]. In this study, we modified the method by using peak height instead of area under the curve (AUC) to estimate production titers. Briefly, diluted samples were loaded onto a C8 column Zorbax 300SB-C8 Rapid Resolution 2.1 × 50 mm, 3.5 μm (Agilent Technologies, Santa Clara, CA) coupled to a C8 guard column Zorbax 300SB-C8, 2.1 × 12.5 mm, 5 μm (Agilent Technologies, Santa Clara, CA) mounted on a Bioinert (Agilent Technologies, Santa Clara, CA). Samples were eluted with 0.1% TFA and 2% n-propanol in LC-MS water as Solvent A, and 0.1% TFA, 70% isopropanol, and 20% acetonitrile in LC-MS water as Solvent B.

### Size Exclusion Chromatography (SEC)

Stored sample supernatants were thawed on ice and buffer-exchanged into 1X PBS pH 7.4 by diafiltration using Amicon® Ultra-15 Centrifugal Filter Units (Millipore, Burlington, MA). The resulting concentrate was loaded into a Superdex® 200 10/300 GL column (Cytiva, Marlborough, MA) and eluted with 1X PBS 500 mM NaCl pH 7.4 at a flow rate of 0.5 mL per minute. The eluted fractions were evaluated against a function test calibration curve using known protein standards to estimate protein sizes.

### Native Gel Electrophoresis

Samples were diluted 1:2 (v/v) with Native Sample Buffer (Bio-Rad, Hercules, CA) and loaded onto 4–20% Mini-PROTEAN® TGX™ Precast Protein Gels (Bio-Rad, Hercules, CA). Electrophoresis was performed at 100 V for 90 min in 1× Tris/Glycine running buffer (Bio-Rad, Hercules, CA) at 4°C.

### Western Blotting

Following gel electrophoresis, proteins were transferred onto a nitrocellulose membrane using the iBlot 2 Dry Blotting System (Invitrogen, Carlsbad, CA) with iBlot^TM^ 2 NC Mini Transfer Stacks (Invitrogen, Carlsbad, CA) following the manufacturer’s transfer protocol. Membrane blocking, primary antibody binding, and secondary labeling were carried out using the iBind^TM^ Western Device (Invitrogen, Carlsbad, CA). Primary broadly neutralizing antibodies (bNAbs) were prepared at final concentrations of 3 μg/mL. Secondary detection was performed using a horseradish peroxidase (HRP)-conjugated goat anti-human IgG (H+L) F(ab’) 2 affinity-purified antibody (1:10,000 dilution; Millipore, Supelco, Burlington, MA). Chemiluminescent signals were captured on a ChemiDoc XRS+ Imaging System (Bio-Rad, Hercules, CA).

### Denaturing Gel Electrophoresis (SDS-PAGE)

Samples were diluted 3:1 (v/v) with 4× Laemmli Sample Buffer (Bio-Rad, Hercules, CA), heated at 95 °C for 5 min, and incubated on ice for 5 min. Following centrifugation at 10,000 × g for 1 min, samples were loaded onto 4–20% Mini-PROTEAN® TGX™ Precast Protein Gels (Bio-Rad, Hercules, CA) and electrophoresed at 100 V for 90 min in 1× Tris/Glycine/SDS running buffer (Bio-Rad, Hercules, CA) at 4°C.

### Coomassie staining

Protein samples were visualized by staining gels with Coomassie Brilliant Blue R-250 (Bio-Rad, Hercules, CA) following standard protocols. Gels were incubated in staining solution for 1 hour at room temperature with gentle agitation, then destained in a methanol-acetic acid solution until the background was clear and protein bands were clearly visible.

### N-Glycan Analysis by MALDI-TOF MS

Samples were processed according to a previously published method with a few modifications [44–46]. Briefly, crude samples were concentrated 15-fold by volume using ultrafiltration centrifugal filters. Concentrated samples were separated via denaturing gel electrophoresis and stained with Coomassie blue as described above. Following destaining, target protein bands were excised and subjected to in-gel glycerol-free PNGase F digestion overnight (New England Biolabs, Ipswich, MA). We recovered the released N-glycans, dried them, and stored them at −20°C. For mass spectrometry, dried glycans were rehydrated and mixed with 2,4,6-trihydroxyacetophenone (THAP) matrix (Millipore Sigma, Supelco, Burlington, MA) for processing of acidic and neutral glycans. MALDI-TOF MS analysis of the N-glycans was performed in reflector positive mode using a Shimadzu MALDI-TOF 7090 mass spectrometer (Shimadzu, Columbia, MD). All samples were analyzed in triplicate under identical experimental conditions. N-glycan structures were assigned by comparing experimental mass values (m/z) against previously published data[44,45].

### Indirect Enzyme-Linked Immunosorbent Assays (ELISA)

Microtiter plates (Sigma-Aldrich, Burlington, MA) were coated overnight at 4°C with 100 µL/well of crude CP-Env (1:50 dilution) or linear gp145 (1:50 dilution) in coating buffer (50 mM Carbonate, pH 9.6). All subsequent steps were conducted at room temperature. Plates were washed three times (200 µL/well, 5 min each) with wash buffer (PBST: 1X PBS with 0.05% v/v Tween-20) and blocked with 200 µL/well blocking buffer (1% w/v non-fat dry whole milk in 1X PBS) for 1 h. After discarding the blocking buffer, primary monoclonal antibodies (3 µg/mL) or diluted patient plasma samples (1:5,000) were added in sample diluent (100 mM HEPES, 100 mM NaCl, 1% w/v BSA, 0.1% v/v Tween-20, pH 7.4;100 µL/well) and incubated for 2 h. Following three washes (200 µL/well, 5 min each), HRP-conjugated secondary antibody (1:5,000 in sample diluent) was added (100 µL/well) for 1 h. Plates were washed three times (200 µL/well, 5 min each) and incubated with 100 µL/well OPD substrate solution (OPD Citrate-Phosphate Buffer, pH 5.0) for 30 min in the dark. Reactions were quenched with 50 µL/well 2.5 M H_2_SO_4_, and absorbance was measured at 490 nm (OD490) within 10 min using a Tecan Infinite M200 Pro reader. Absorbance values (OD490) were background-corrected by subtracting the mean optical density of blank wells and plotted in GraphPad Prism version 10.3.1 (San Diego, CA). Plasma immunoreactivity cutoffs were defined as the mean OD490 of PLWOH plasma samples plus 3 standard deviations (3 x SD). Differences among groups were assessed using one-way ANOVA followed by Tukey’s post-hoc test for multiple comparisons. Statistical significance was set at *p* < 0.05.

### Antibodies and Reference Controls

The broadly neutralizing antibodies (bNAbs) used in this study were obtained from the NIH HIV Reagent Program, Division of AIDS, NIAID, NIH (now available through BEI Resources; formerly the NIH AIDS Reagent Program). These include monoclonal anti-HIV-1 gp120 (VRC01, produced *in vitro*, Cat. No. ARP-12033), monoclonal anti-HIV-1 gp120 (PG9, Cat. No. ARP-12149) contributed by the International AIDS Vaccine Initiative, monoclonal anti-HIV-1 gp120 (PG16, Cat. No. ARP-12150), monoclonal anti-HIV-1 gp120 (PGT121, Cat. No. ARP-12343) contributed by the International AIDS Vaccine Initiative, monoclonal anti-HIV-1 gp120 (2G12, Cat. No. ARP-1476), and monoclonal anti-HIV-1 gp120 (4E10, Cat. No. ARP-10091) [47–50]. The secondary antibody used was goat anti-human IgG (H+L) F(ab’)2 HRP-conjugated (Millipore, Burlington, MA).

Plasma samples from four people living without HIV (PLWOH) were used as controls (n=4), and ten plasma samples (n=10) from people living with HIV (PLWH) were generously provided by the Repository of the NeuroHIV Clinical Cohort of Adults with HIV infection (IRB protocol 2290032835R004, Cantres-Rosario et al., 2022; Wojna et al., 2006). **Table 2** describes participant characteristics.

The recombinant Linear-C06980v0c22-K3 used as a linear control was kindly provided by CDI Laboratories (Puerto Rico, USA). Linear-C06980v0c22-K3 used for western blot analysis was obtained as a purified protein via sequential HiTrap Capto Lentil Lectin and Toyopearl NH2-750F chromatography.

## Results

### Design strategy and production of circularly permuted Env

We hypothesized that circularly permuting truncated Env sequences would yield stable trimers capable of binding broadly neutralizing antibodies (bNAbs). To test this strategy, we engineered four circularly permuted (CP) constructs using the BG505 (GenBank: ABA61515) and C06980v0c22 [41] *env* genes as templates. For CP-BG505, we incorporated the T332N mutation to enable bNAb recognition of the glycan supersite [46,53,54]. We also inserted flexible linkers (6- and 20-amino acid) between the C-terminus of gp41 and the N-terminus of gp120 to facilitate trimerization [38]. For CP-C06980v0c22-K3, instead of linkers, we added three lysine residues (K3) to the C-terminus of gp41 to enhance expression and reduce aggregation [42]. CP constructs were expressed in FreeStyle CHO-S cells and harvested after nine days in culture. Production yields were determined using a previously described analytical reverse-phase HPLC method (RP-HPLC; González-Feliciano et al., 2020). As shown in **Fig. 2A**, CP-Env eluted at 5.29 minutes, and peak height was used for quantification. All CP-Env constructs were successfully expressed, with yields averaging 48.15 ± 6.62 mg/L (**Fig. 2B**).

**Fig. 2.**
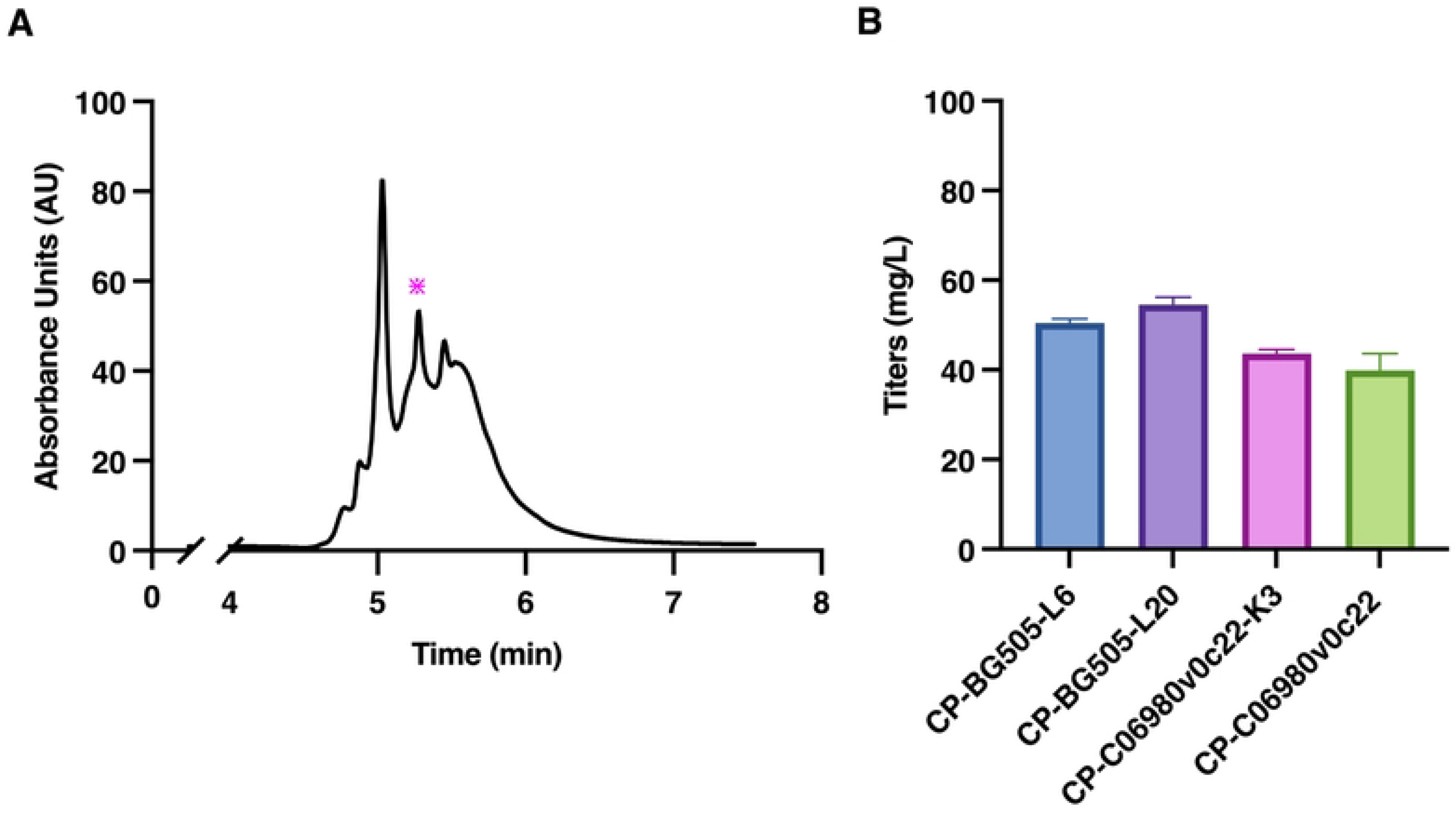
RP-HPLC analysis and production yields of recombinant Env variants in FreeStyle CHO-S cells. (A) Representative analytical RP-HPLC chromatogram showing the retention time and intensity (A_280_) of the secreted proteins. The asterisk (*) denotes the elution fraction containing CP-C06980v0c22-K3, confirmed by immunoblot (data not shown). (B) Volumetric harvest titers (mg/L) of recombinant Env constructs after 9 days in culture. Bars represent mean ± SD (n=3).

### Oligomeric state of circularly permuted Env

The oligomeric state of CP-C06980v0c22 was evaluated by size-exclusion chromatography (SEC) and western blotting. The SEC chromatogram revealed six partially resolved peaks (**Fig. 3A**). Native western blotting confirmed the presence of CP-C06980v0c22 in fractions 2 and 3 through recognition by the broadly neutralizing antibody 4E10 (**Fig. 3B**). The retention time for Fraction 2 (R_t_ = 21.06 min) corresponded to an apparent molecular weight of 397 kDa, approximately 2.7 times the theoretical monomer mass (145 kDa), indicating a possible trimeric assembly. Western blot analysis revealed a mixture of higher-order aggregates and trimers, with trimers representing the predominant species (**Fig. 3B**). The retention time for Fraction 3 (R_t_ = 23.34 min) corresponded to an apparent molecular weight of 229.3 kDa, consistent with a predominantly dimeric assembly. The western blot showed a mixture of both trimeric and dimeric species (**Fig. 3B**).

**Fig. 3.**
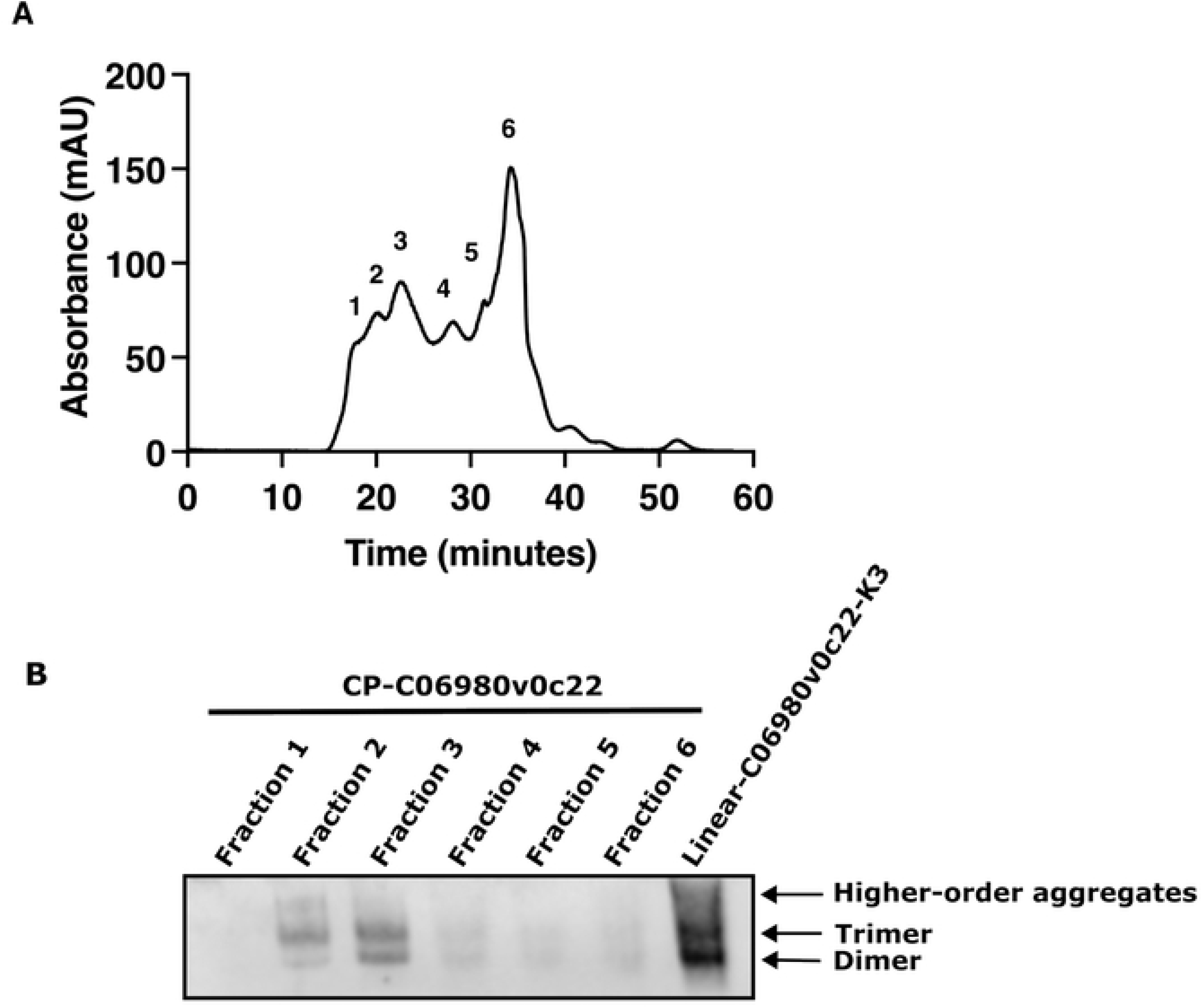
Size-exclusion chromatography (SEC) profile and Native-PAGE western blot analysis of CP-C06980v0c22. (A) Preparative SEC elution profile and oligomeric state estimation: SEC chromatogram monitoring UV absorbance at 280 nm (A_280_) as a function of retention time to evaluate the quaternary species distribution of the recombinant Env construct. Numbered peaks (Fractions 1–6) indicate the collected fraction windows. Apparent molecular weights (M_r_) were calculated relative to calibration standards to assign oligomeric states, resolving higher-order aggregates, native-like trimers, and dimers. (B) Native-PAGE western blot of SEC fractions: Western blot of a non-denaturing 4–10% acrylamide gradient Native-PAGE gel probed with the MPER-directed broadly neutralizing antibody 4E10. The preservation of native quaternary assemblies confirms the structural integrity and oligomeric assignments of the SEC fractions from (A). Lanes 1–6: collected SEC fractions 1–6; Lane 7: reference control (purified Linear-C06980v0c22-K3 with defined oligomeric distribution).

### Glycosylation profile of circularly permuted Env

We profiled PNGase F-released *N*-linked glycans by MALDI-TOF MS. Comparative analysis revealed broad similarities between glycans released from Linear-C06980v0c22-K3 (**Fig. 4A**) and CP-C06980v0c22 (**Fig. 4B**). Both profiles were dominated by high-mannose structures (Man5–Man9), consistent with those previously obtained for Linear-C06980v0c22-K3 [45]. However, CP-C06980v0c22 showed greater structural diversity due to additional complex-type glycans (**Table 1**).

**Fig. 4.**
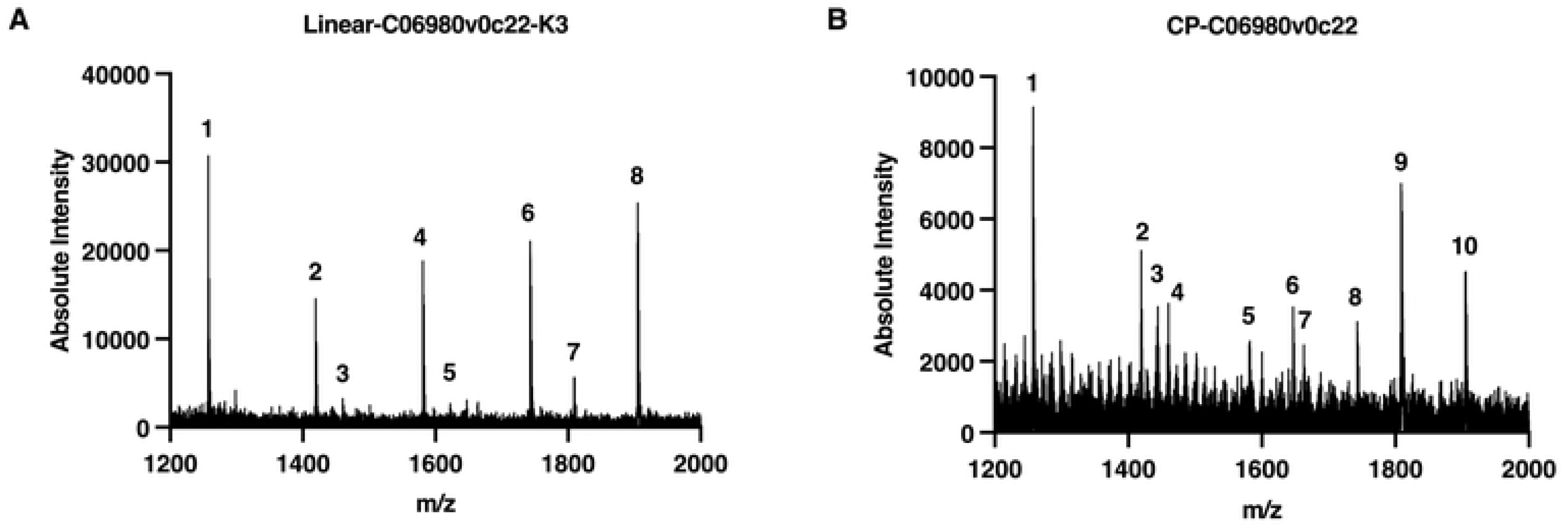
MALDI-TOF MS profiling of PNGase F-released N-linked glycans from Linear-C06980v0c22-K3 (A) and CP-C06980v0c22 (B). Representative MALDI-TOF mass spectrum of *N*-linked glycans enzymatically released from recombinant Env constructs using PNGase F and acquired in positive ion mode across a mass-to-charge ratio (*m/z)* range of 1,200 – 2,000. Numbered peaks correspond to the specific glycan compositions and assignments summarized by Peak ID in **Table 1**. All observed glycan species are detected as sodium adducts ([M + Na]^+^).

**Table 1.**
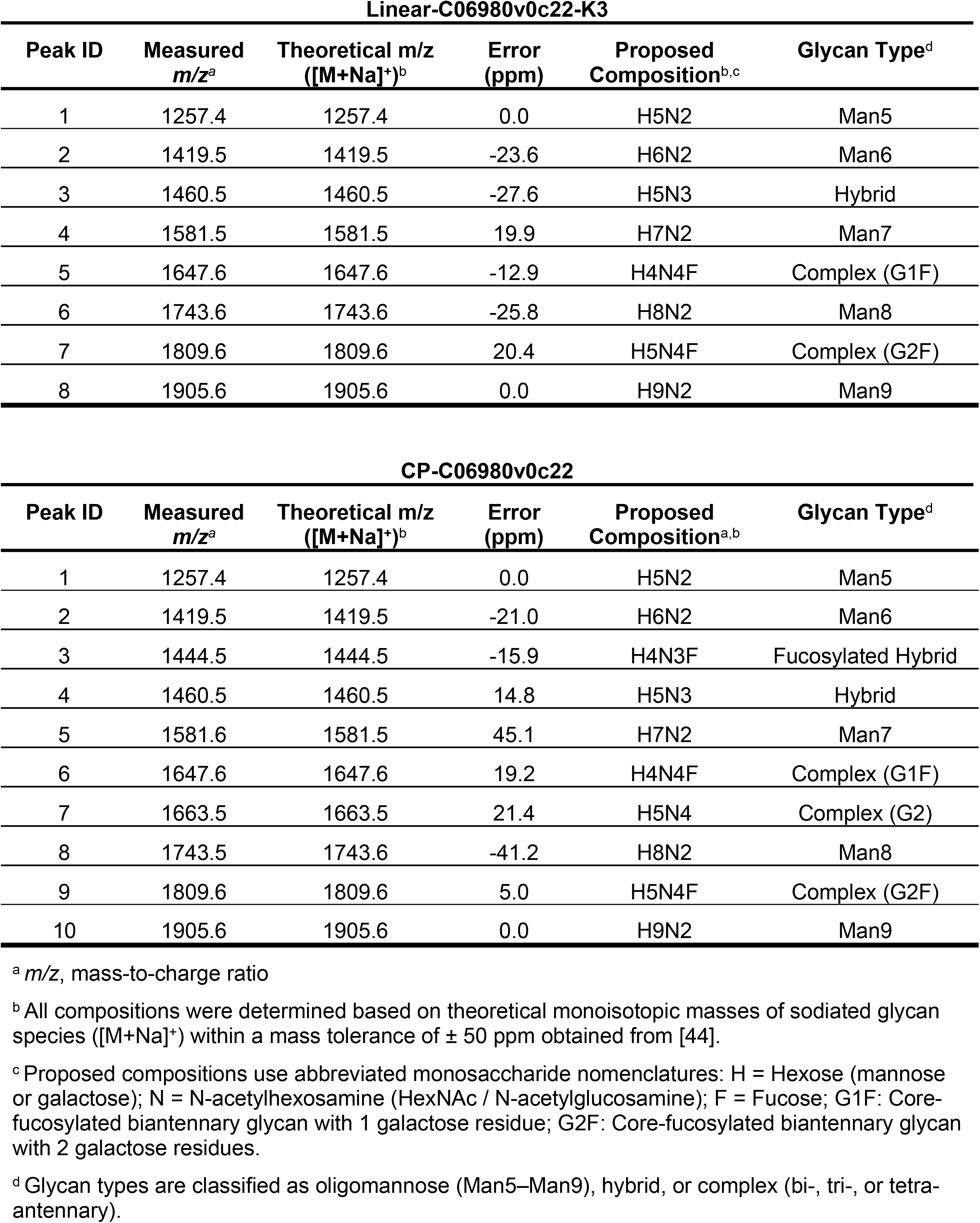
Mass spectrometric assignment of PNGase F-released *N*-linked glycans from Linear-C06980v0c22-K3 and CP-C06980v0c22. Summary of peak assignments from positive-mode MALDI-TOF MS spectra (**Fig. 4**). Peak IDs correspond to the numbered annotations in **Fig. 4**.

| Linear-C06980v0c22-K3 |  |  |  |  |  |
| Peak ID | Measured<br>*m/z<sup>a</sup>* | Theoretical m/z<br>([M+Na]<sup>+</sup>)<sup>b</sup> | Error<br>(ppm) | Proposed<br>Composition<sup>b,c</sup> | Glycan Type<sup>d</sup> |
|---|---|---|---|---|---|
| 1 | 1257.4 | 1257.4 | 0.0 | H5N2 | Man5 |
| 2 | 1419.5 | 1419.5 | -23.6 | H6N2 | Man6 |
| 3 | 1460.5 | 1460.5 | -27.6 | H5N3 | Hybrid |
| 4 | 1581.5 | 1581.5 | 19.9 | H7N2 | Man7 |
| 5 | 1647.6 | 1647.6 | -12.9 | H4N4F | Complex (G1F) |
| 6 | 1743.6 | 1743.6 | -25.8 | H8N2 | Man8 |
| 7 | 1809.6 | 1809.6 | 20.4 | H5N4F | Complex (G2F) |
| 8 | 1905.6 | 1905.6 | 0.0 | H9N2 | Man9 |
| CP-C06980v0c22 |  |  |  |  |  |
| Peak ID | Measured<br>*m/z<sup>a</sup>* | Theoretical m/z<br>([M+Na]<sup>+</sup>)<sup>b</sup> | Error<br>(ppm) | Proposed<br>Composition<sup>a,b</sup> | Glycan Type<sup>d</sup> |
| 1 | 1257.4 | 1257.4 | 0.0 | H5N2 | Man5 |
| 2 | 1419.5 | 1419.5 | -21.0 | H6N2 | Man6 |
| 3 | 1444.5 | 1444.5 | -15.9 | H4N3F | Fucosylated Hybrid |
| 4 | 1460.5 | 1460.5 | 14.8 | H5N3 | Hybrid |
| 5 | 1581.6 | 1581.5 | 45.1 | H7N2 | Man7 |
| 6 | 1647.6 | 1647.6 | 19.2 | H4N4F | Complex (G1F) |
| 7 | 1663.5 | 1663.5 | 21.4 | H5N4 | Complex (G2) |
| 8 | 1743.5 | 1743.6 | -41.2 | H8N2 | Man8 |
| 9 | 1809.6 | 1809.6 | 5.0 | H5N4F | Complex (G2F) |
| 10 | 1905.6 | 1905.6 | 0.0 | H9N2 | Man9 |

<sup>a</sup> *m/z*, mass-to-charge ratio
<sup>b</sup> All compositions were determined based on theoretical monoisotopic masses of sodiated glycan species ([M+Na]<sup>+</sup>) within a mass tolerance of ± 50 ppm obtained from [44].
<sup>c</sup> Proposed compositions use abbreviated monosaccharide nomenclatures: H = Hexose (mannose or galactose); N = N-acetylhexosamine (HexNAc / N-acetylglucosamine); F = Fucose; G1F: Core-fucosylated biantennary glycan with 1 galactose residue; G2F: Core-fucosylated biantennary glycan with 2 galactose residues.
<sup>d</sup> Glycan types are classified as oligomannose (Man5–Man9), hybrid, or complex (bi-, tri-, or tetra-antennary).

**Table 2.** Participant characteristics. Summary of cohort characteristics from people living without HIV (PLWOH) and people living with HIV (PLWH). The clinical data was accessed on May 20, 2026.

| Parameter | PLWOH (n = 4) | PLWH (n = 10) |
|---|---|---|
| **Age** (years<sup>1</sup>) | 27 (25, 29) | 39 (24, 58) |
| **CD4 cells/mm<sup>3</sup>** | -- | 395.5 (61, 1007) |
| **CD8 cells/mm<sup>3</sup>** | -- | 663.5 (416, 1246) |
| **CD4/CD8 ratio** | -- | 0.531 (0.125, 1.459) |
| **CD4 Nadir** (cells/mm<sup>3</sup>) | -- | 150 (40, 331) |
| **HIV RNA copies/mL** (log<sub>10</sub>) | -- | 2.10 (1.30, 5.50) [4 ND<sup>2</sup>, 2 missing] |
| **Body Mass Index** (kg/m<sup>2</sup>) | 35.4 (1 missing) | 28.5 (22.5, 41.0) |
| **ART<sup>3</sup> combinations** (n = 3) | -- | zidovudine, lamivudine, nelfinavir, tenofovir DF, abacavir *(Combivir, Viracept, Viread, Epzicom)* |
| **new ART<sup>3</sup> combinations** (n = 3) | -- | raltegravir, atazanavir, etravirine, emtricitabine, tenofovir DF *(Isentress, Reyataz, Intelence, Truvada)* |
| **Unspecified / No ART** | -- | 1 unspecified, 3 none |
| **Neurocognitive Status (HAND<sup>4</sup>)** | -- | 3 Normal, 2 ANI, 2 MND, 1 HAD (2 missing) |
| **HCV Status (Positive)** | -- | 1 / 9 (11.1%) [1 missing] |
| **Toxicology (Cocaine Positive)** | -- | 2 / 10 (20.0%) |

<sup>1</sup> Median (range: min, max)
<sup>2</sup> ND = Not detectable (detection limit: 50 copies/mL for old regimens/log 1.70; 20 copies/mL for new regimens/log 1.30)
<sup>3</sup> ART = Antiretroviral treatment
<sup>4</sup> HAND = HIV-Associated Neurocognitive Disorder (ANI: Asymptomatic Neurocognitive Impairment; MND: Mild Neurocognitive Disorder; HAD: HIV-Associated Dementia)

### Recognition of circularly permuted Env by broadly neutralizing antibodies

To determine whether the CP constructs displayed native-like epitopes, we evaluated their recognition by a panel of six bNAbs (PG16, PG9, PGT121, 2G12, 4E10, and VRC01) using western blot analysis under native and denaturing conditions (**Fig. 5**). Linear-C06980v0c22-K3 served as a positive control due to its strong recognition by the bNAb panel [31]. Because its initial signal obscured the CP construct bands and prevented resolution of individual oligomeric states, we loaded the linear control at an 8-fold lower concentration. Under native conditions, the bNAb panel demonstrated distinct construct preferences and oligomeric resolution profiles. PG16, PG9, 4E10, and VRC01 exhibited stronger reactivity toward CP-C06980v0c22-K3 and CP-C06980v0c22, than to the BG505 versions. In contrast, PGT121 preferentially bound CP-BG505-L6 and CP-BG505-L20, while 2G12 bound exclusively CP-BG505-L6 and CP-BG505-L20 and showed no detection of the linear control. PG16 and PG9 primarily recognized trimers and higher-order aggregates. PGT121, 2G12, 4E10, and VRC01 detected three species: dimers, trimers, and higher-order aggregates. Denaturing conditions severely restricted bNAb recognition of CP constructs. PG9, 2G12, 4E10, and VRC01 retained binding in the denatured state, recognizing CP-C06980v0c22-K3, CP-C06980v0c22, and Linear-C06980v0c22-K3. CP-BG505-L6 and CP-BG505-L20 were not recognized under denaturing conditions.

**Fig. 5.**
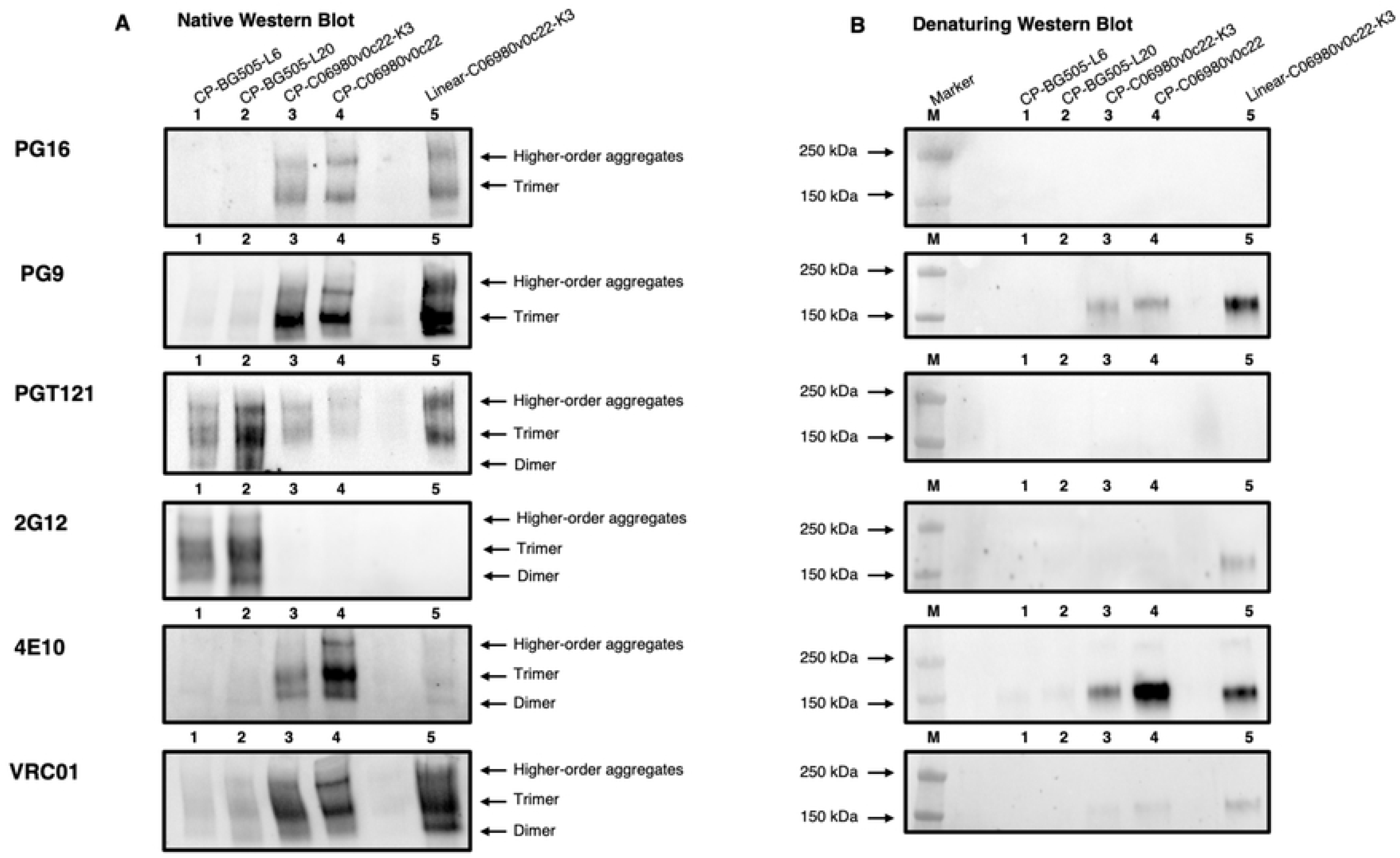
Native-PAGE and denaturing SDS-PAGE western blot characterization of CP Env variants. (A) Native-PAGE western blot (quaternary structure profile): Non-denaturing Native-PAGE western blot of CP Env variants probed with six broadly neutralizing antibodies (bNAbs: PG16, PG9, PGT121, 2G12, 4E10, and VRC01) targeting distinct Env domains (V1/V2 apex, V3/glycan patch, MPER, and CD4-binding site). Arrows denote distinct oligomeric species, including higher-order aggregates, trimers, and dimers. Lanes 1–5: Env constructs CP-BG505-L6, CP-BG505-L20, CP-C06980v0c22-K3, CP-C06980v0c22, and Linear-C06980v0c22-K3, respectively. (B) Reducing SDS-PAGE western blot (subunit integrity): Denaturing, reducing (with 2-mercaptoethanol) SDS-PAGE western blot of the corresponding Env variants probed with the same bNAb panel to assess individual subunit molecular weight and primary polypeptide integrity under reducing conditions. Lane M: molecular weight marker (kDa); Lanes 1–5: Env constructs CP-BG505-L6, CP-BG505-L20, CP-C06980v0c22-K3, CP-C06980v0c22, and Linear-C06980v0c22-K3, respectively.

### Immunoreactivity of circularly permuted Env against patient-derived antibodies

To further evaluate if CP constructs present native-like epitopes, we performed a seroreactivity assay using patient plasma. The reactivity of CP and linear constructs was tested against plasma samples from ten people living with HIV (PLWH, n=10) and four people living without HIV (PLWOH, n=4). As expected, the seroreactivity of all constructs was significantly higher with plasma from PLWH than with PLWOH (*p* < 0.05; **Fig. 6A**). All CP constructs exhibited immunoreactivity above the cutoff threshold (mean + 3 SD). Yet, they were less reactive than Linear-C06980v0c22-K3, likely reflecting differences in key epitope exposure (*p* < 0.05; **Fig. 6B**).

**Fig. 6.**
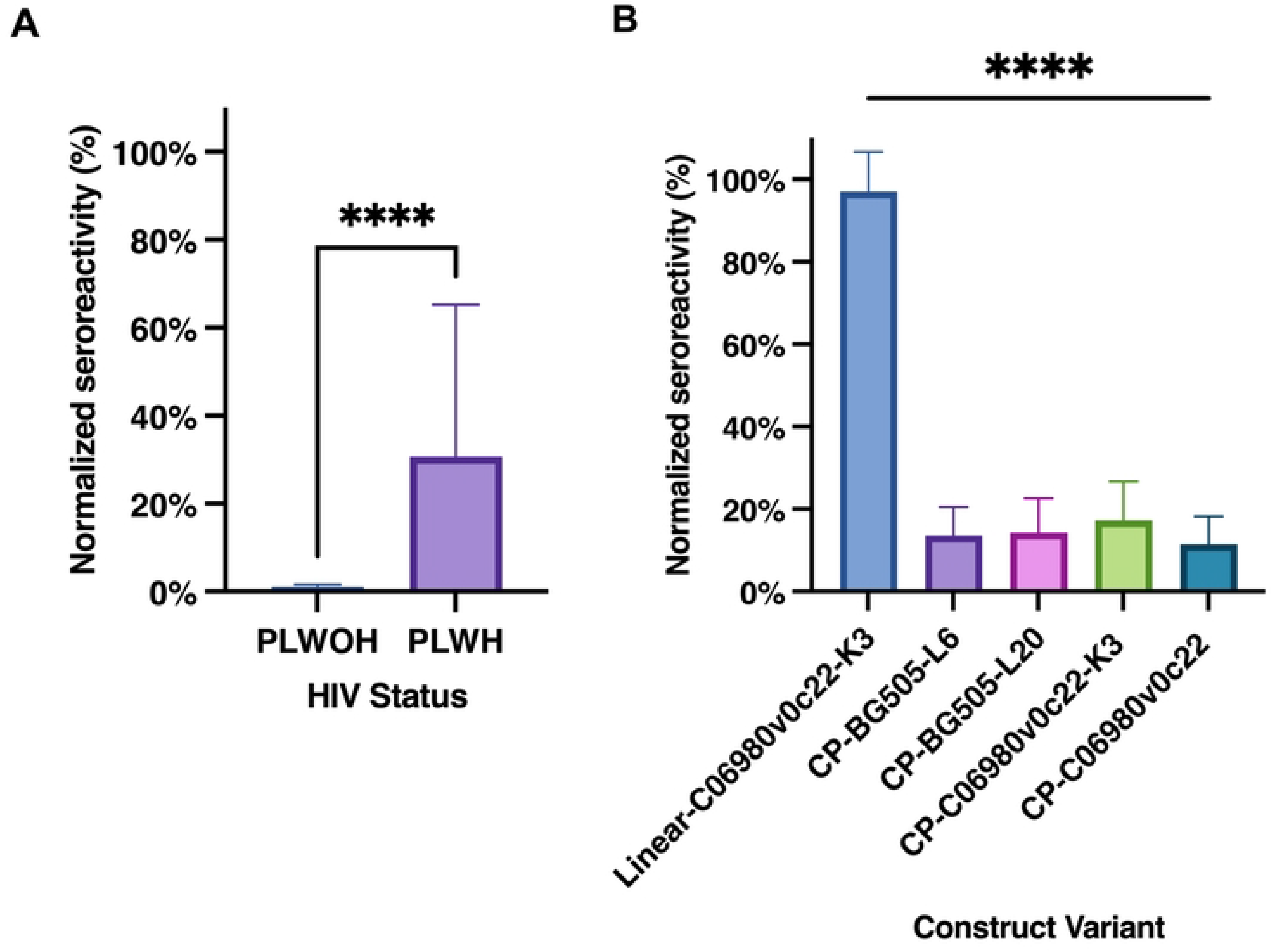
Comparative seroreactivity of recombinant Env constructs across clinical cohorts and structural formats. (A) Cohort seroreactivity: Relative plasma antibody binding (normalized seroreactivity, %) in PLWOH (n = 4) compared to PLWH (n = 10). Seroreactivity values are normalized relative to PLWH 1 (set to 100%). PLWH exhibited significantly higher seroreactivity than PLWOH (p<0.05, two-tailed Student’s *t*-test). (B) Structural format seroreactivity: Normalized seroreactivity (%) of CP Env constructs compared to Linear-C06980v0c22-K3. Values are normalized relative to PLWH 1 (set to 100%). CP constructs displayed significantly reduced seroreactivity relative to the linear antigen (p<0.05, one-way ANOVA with Tukey’s post hoc test). Data are presented as mean + SD, with statistical significance (p<0.05) indicated by an asterisk (*).

## Discussion

Despite years of vaccine research, achieving sustained immunity against HIV-1 at population level remains a major challenge. Numerous HIV vaccine trials have explored a range of immunization strategies that incorporate some form of the HIV-1 Env protein. Unlike internal HIV proteins, including Gag, Pol, or Nef, Env is the only viral protein exposed on the surface of free intact virions [55] and is the only one capable of eliciting broadly neutralizing antibodies that prevent viral entry into host cells [56–58]. To optimize Env presentation, trial strategies have developed a range of platforms, including virus-like particles [59–61], recombinant viral vectors [62–64], nanoparticle displays [6,65], and soluble recombinant proteins [20]. Unfortunately, these protein design and delivery strategies face challenges like antigen structural instability, exposure of distracting epitopes, off-target responses, and anti-vector immunity. Thus, immunological strategies that provide broad-spectrum protection and support diverse populations remain a global aspiration that has not yet been achieved.

In this work, we report the production and characterization of circular permutant constructs of Env gp145 featuring an N-terminal gp41 followed by a C-terminal gp120. The resulting constructs display core antigenic and structural features of uncleaved genetically fused trimers. CHO cells successfully produced these constructs at detectable levels in culture medium, as confirmed by western blotting and RP-HPLC. Production levels were comparable to those achieved in other efforts, exceeding 40 mg/L [66]. Therefore, optimizing the cell line and expression strategy could yield even higher production levels, thereby improving the feasibility of production for clinical trials [67–69].

CP Envs produce an oligomeric mixture that is predominantly trimeric, as observed by SEC with an estimated molecular weight of 397 kDa. This aligns with analytical studies showing that BG505 DS-SOSIP.664 Env trimers can range from 342 to 316 kDa [69]. The oligomeric state was also confirmed by native western blots using the bNAbs PG16 and PG9, which recognize trimeric species [70]. Maintaining this trimeric identity is important because it more closely resembles the native form. Previous reports show that native-like HIV Env trimers have significantly greater neutralization breadth and potency than monomeric subunits [29,71].

The circular permutants in this study were recognized by the glycosylation machinery of the cells, resulting in patterns that resemble the glycosylation of Env in linear native trimers, as demonstrated by MALDI-TOF MS analysis. However, the CPs displayed a higher proportion of complex-type glycans compared to their cognate linear construct. This enrichment in complex glycans departs from the typical oligomannosidic pattern, as complex glycans are larger and more branched than mannose species. Conversely, this shift in glycosylation might expose more protective epitopes and shield immunodominant, non-neutralizing epitopes [72].

Because Env antigenicity is a critical benchmark for immunogen evaluation, we assessed how circular permutation affects binding recognition across a panel of bNAbs targeting major sites of vulnerability. Remarkably, these bNAbs demonstrated significant selectivity across strains and construct variants. These results were unexpected, given that we had selected a panel of bNAbs known for their high affinity for previously characterized trimers from the BG505 and C06980 strains [22,31,45,73]. To evaluate epitope exposure and structural integrity, we compared antibody reactivity under both native and denaturing western blot conditions. A subset of bNAbs showed clear preference for the C06980 strain over BG505, including PG16, PG9, 4E10, and VRC01. Both PG16 and PG9 belong to the family of V2-apex glycan-dependent trimer antibodies. Studies of the BG505 SOSIP.664 gp140 reveal that these bNAbs recognize the trimer asymmetrically through contacts with two gp120 protomers and conserved glycans at positions N156 and N160 [74,75], with PG9 displaying higher affinity [76,77]. PG16 reactivity was observed exclusively in the native state and was absent under denaturing conditions, confirming that both CP and linear constructs of C06980 strains form native-like, quaternary-assembled apex structures. In contrast, PG9 bound all constructs under both native and denaturing conditions, preserving reactivity under denaturing conditions. This indicates that PG9 can bind sequence features that remain after unfolding, whereas PG16 strictly requires quaternary assembly [78].

The bNAbs 4E10 and VRC01 target distinctive functional domains on different subunits of Env. Recognition of Env by 4E10 is dependent on conserved residues W672, F673, T676, and W680 of the MPER of gp41 [78]. The strong reactivity of 4E10 under both native and denaturing Western blot conditions indicates greater exposure and accessibility of the MPER epitope in the CP-C06980 variants than in the linear construct. In contrast, VRC01 binds the conformationally sensitive CD4-binding site (CD4bs) on gp120 with broad clade coverage [49,79,80]. Strong VRC01 binding under native conditions demonstrates proper structural assembly and exposure of the CD4bs of these constructs.

Conversely, the bNAbs 2G12 and PGT121 preferentially bound BG505 over C06980, engaging the N332 glycan-dependent clusters present on both constructs. Although both antibodies depend on N332, they represent distinct structural classes. 2G12 targets purely mannose-dependent clusters on the outer domain of gp120, whereas PGT121 recognizes a composite peptide–glycan epitope at the base of the V3 loop that accommodates complex-type glycans [22,81–84]. Crucially, PGT121 bound strongly under native conditions but produced no signal on denaturing western blots, confirming that both CP and linear constructs of C06980 strains display native-like V3 base architectures. In contrast, the reduced binding of 2G12 to C06980 relative to BG505 suggests that outer domain glycans in C06980 are less accessible or structurally altered.

Finally, we tested the seroreactivity of our CP constructs relative to their linear cognates using polyclonal plasma from PLWOH and PLWH. Our participant cohort consisted of individuals from Puerto Rico, where clade B accounts for 90% of infections [85], a different lineage from our clade A (BG505) and Clade C (C06980v0c22) immunogens. Due to this heterologous clade mismatch, overall reactivity was expected to be low.

Nevertheless, all constructs were more immunoreactive against plasma from PLWH than from PLWOH controls. Notably, all circularly permuted C06980 constructs were significantly less reactive than the linear cognates. This decreased seroreactivity suggests that circular permutation induces structural alterations that either shield a subset of conserved epitopes required for broad cross-reactivity or mask immuno-distracting non-neutralizing epitopes. Taken together, these results provide a framework to further refine antigen presentation in future immunogen engineering.

Overall, these findings highlight the clinical relevance of circular permutants as a promising platform for the rational design of HIV immunogens. While current evaluations were limited to clades A and C in a local cohort, future efforts should expand the platform’s translational scope to additional clades across epidemiologically diverse populations.

## Conclusions

As the search for improved vaccine platforms advances, circular permutation offers a promising strategy for engineered immunogen design. Our results show that this approach preserves key native Env characteristics, such as trimerization, epitope presentation, and glycosylation profiles, while masking non-neutralizing regions. Moving forward, prioritizing immunogen stability and targeted immunoreactivity will remain essential for these platforms to guide future vaccine development.

## Acknowledgements

The authors thank Ms. Vilmarie Mercado for administrative support; Dr. José González-Feliciano for valuable technical guidance across cell culture, protein purification, and MALDI-TOF MS; and Dr. Rafael Maldonado-Hernández for guidance on protein purification strategies. We also thank Dr. Juan Villalobos for key insights regarding cell culture, Mr. Ohel Soto for cell culture maintenance, Mr. Bryan Rodriguez for assisting with sample purification, and Mr. Matthew Trueblood and Mr. Juan Pacheco for handling MALDI-TOF MS samples. We also thank Ms. Gabriela Acevedo, Dr. Dina Bracho, and Dr. Coral Capó for their guidance with ELISA assays.

